# Hybrid Stacking–Bagging Ensembles for Robust Multi-Omics Breast Cancer Prognosis

**DOI:** 10.64898/2026.01.21.700935

**Authors:** Reza Bozorgpour

## Abstract

Accurate breast cancer risk prediction remains a central challenge in precision oncology due to the complexity and heterogeneity of underlying biological processes. While single-modality models based on clinical, gene expression, or copy number variation (CNV) data provide valuable prognostic insights, they often fail to capture complementary information across data sources. Conventional stacking ensembles improve predictive performance through multimodal integration but remain susceptible to variance and overfitting.

In this study, we propose a heterogeneous hybrid ensemble framework that combines stacking and bagging to enhance robustness and accuracy in multi-omics breast cancer classification. The framework integrates clinical features, gene expression profiles, and CNV data through stacked multimodal representations, followed by parallel stacking and bagging meta-learning and weighted fusion. Experiments conducted on the METABRIC cohort demonstrate that the proposed hybrid model achieves a ROC AUC of 0.9355, outperforming unimodal models (AUC range: 0.80–0.88) and a conventional stacking ensemble (AUC = 0.919). At the Youden’s J optimal operating point, the hybrid approach yields balanced sensitivity (0.8571) and specificity (0.8792), with an overall accuracy of 87.4% and an F1-score of 0.7706.

These results highlight the effectiveness of hybrid ensemble learning for robust multimodal integration and demonstrate its potential as a scalable and reliable approach for breast cancer risk prediction. The proposed framework offers a practical pathway toward improved predictive stability and supports the broader application of ensemble-based strategies in precision medicine.

## Introduction

Breast cancer is a major cause of global mortality, predominantly affecting women [1-3]. It results from abnormal breast cell growth, producing benign tumors that remain localized or malignant tumors that can invade and metastasize. The disease is classified as invasive or non-invasive, depending on whether cancer cells spread beyond the ducts or lobules into surrounding tissue [4, 5]. Although far less common, breast cancer can also occur in men [6]. In 2020, approximately 2.26 million new cases were reported worldwide, and in the United States an estimated 287,850 invasive and 51,400 non-invasive cases were diagnosed, with about 43,580 deaths projected in 2022. Breast cancer prognosis is typically defined as short-term (<5 years) or long-term (>5 years) survival [4]. Survival rates vary substantially by disease stage: non-invasive cancer shows 90% five-year and 84% ten-year survival, while invasive cases range from 99% when localized to 86% with lymph node involvement and 29% with distant metastasis. This heterogeneity makes accurate prognosis difficult but critical for individualized treatment planning [7]. Advances in machine learning and deep learning have transformed medical diagnostics and prognosis [8, 9]. Deep learning can automatically extract predictive features and has demonstrated the ability to identify cancer prior to clinical manifestation [10, 11]. Prognostic performance can be further improved through hyper-parameter optimization and multimodal data integration, including clinical, gene expression, and CNV data; however, most existing methods rely on homogeneous deep learning models across modalities, limiting their effectiveness [12].

Advances in gene expression profiling have enabled the identification of molecular signatures for breast cancer prognosis, including early gene-based predictors of metastasis in lymph node–negative patients [13]. As breast cancer is fundamentally genetic, subsequent studies incorporated multimodal data—such as clinical variables, gene expression, and CNV—to improve prognostic accuracy and capture complex biological interactions [14]. Kernel-based and machine learning approaches, including GPMKL and molecular prognostic scoring systems, demonstrated improved prediction performance but were limited by data scale or feature selection constraints [15, 16]. Recent deep learning methods address these limitations through multimodal integration. Dense neural networks, autoencoder-based fusion models, and convolutional architectures have shown improved performance in survival prediction and multi-class classification tasks [17-19]. Feature selection frameworks further clarified relationships between prognosis and prediction [20], while mathematical modeling offered complementary insights into tumor growth and treatment response [21]. Finally, scalable multimodal survival models such as MultiSurv extended these approaches across multiple cancer types and data modalities, demonstrating robust long-term survival prediction even with missing data [22].

Ensemble and multimodal deep learning models have become increasingly important for breast cancer diagnosis and prognosis. Several studies ensemble CNN architectures and optimization techniques for imaging-based detection, including ultrasound-based feature extraction using VGG and SqueezeNet models [23], MRI-driven survival and treatment response prediction via stacking and integration strategies [24], and hybrid transfer learning with 2D/3D mammograms achieving improved classification accuracy [25]. Optimization-based classifiers combining WOA, DA, and RBF-SVM have also shown strong performance on benchmark datasets [26].

Beyond imaging, multimodal survival prediction frameworks integrate genomic and clinical data. Graph-based multimodal learning using gene expression, CNV, and pathology data has achieved high accuracy and AUC for survival classification [27], while snapshot ensemble neural networks with t-SNE-based feature reduction improved diagnostic performance [28]. Adversarial multimodal representation learning combining clinical and imaging data further enhanced prognosis prediction [29]. Stacked ensemble approaches using CNN feature extraction followed by Random Forest classification demonstrated strong survivability prediction using multimodal inputs, and weighted fusion frameworks integrating imaging and gene data achieved reliable subtype classification [30].

Earlier prognosis models relied on a single modality, such as gene expression alone [31], which limited predictive capability. To address this, multimodal deep learning architectures integrating clinical data, CNV, and gene expression were introduced, including Multimodal Deep Neural Networks (MDNNMD) [18] and dual-stage stacked ensemble models based on homogeneous CNNs]. However, prior studies indicate that heterogeneous ensemble architecture provides improved generalization over homogeneous designs [32]. Motivated by these findings, the proposed work introduces a heterogeneous multimodal hybrid ensemble framework that leverages stacking for multimodal feature integration and bagging for variance reduction, integrating clinical, gene expression, and CNV data to improve robustness and accuracy in breast cancer risk prediction.

## Materials and methods

### Dataset

This study utilizes the publicly available METABRIC breast cancer cohort, which can be accessed at (https://www.cbioportal.org/study/summary?id=brca_metabric). The dataset contains 1,980 breast cancer patients with complete records and has been widely used for survival analysis and prognostic modeling [33]. For each patient, the dataset provides multi-modal information, including clinical variables, gene expression profiles, and CNV data. Patients were stratified into two outcome groups based on 5-year overall survival. Individuals who survived longer than five years following diagnosis were assigned to the long-term survival group, whereas those who died within five years were categorized as short-term survivors. This stratification resulted in 1,489 long-term survivors and 491 short-term survivors. The median age at diagnosis across the cohort is 61 years, with an average survival duration of 125.1 months. For modeling purposes, long-term survivors were encoded as class ‘0’, while short-term survivors were encoded as class ‘1’. A summary of the dataset characteristics is provided in Table 1. Among the 1,980 patients, 64 individuals (3.23%) lacked complete five-year follow-up information. Consistent with prior studies, these cases were treated as long-term survivors in our analysis.

**Table 1:** Overall METABRIC dataset information.

|  |  |
| --- | --- |
| **Renounce Years** | 5 |
| **Number of patients** | 1980 |
| **Number of patients survived more than 5 years** | 1489 |
| **Number of patients survived less than 5 years** | 491 |
| **Diagnosis median age** | 61 |
| **Average survival month** | 125.1 |

### Data Preprocessing

The METABRIC dataset integrates clinical and molecular information from breast cancer patients. Clinical variables were obtained and linked to tumor molecular features by matching patient identifiers to corresponding sample identifiers. This integration resulted in a unified dataset containing both patient-level clinical characteristics and tumor-specific molecular attributes, yielding a total of 34 variables. The combined dataset was used for all further preprocessing. Data preprocessing was performed to improve consistency and reduce redundancy. The *3-Gene classifier subtype* variable contained categories that deviated from the standard three-gene framework by embedding proliferation information; therefore, proliferation status was extracted into a separate variable, and subtype labels were reassigned based on ER, PR, and HER2 expression. The *Sex* variable exhibited no variability and was removed. Redundancy was identified between *Tumor Other Histologic Subtype* and *Oncotree Code*, with a many-to-one relationship in which multiple histologic subtypes mapped to a single Oncotree classification. In particular, the *IDC* code corresponded to three distinct subtypes, while all other codes showed one-to-one correspondence. Missing histological subtype values were recoverable from *Oncotree Code*. Accordingly, *IDC* was refined into its corresponding subtypes, and *Tumor Other Histologic Subtype* was excluded. A one-to-one relationship was also observed between *Oncotree Code* and *Cancer Type Detailed*, leading to removal of the latter. Finally, *Patient ID, ER status measured by IHC, HER2 status measured by SNP6*, and *Cohort* were removed, as these variables were either non-informative identifiers or redundant with harmonized receptor status fields.

The *Cancer Type* variable was removed because nearly all patients were diagnosed with breast cancer, with breast sarcoma observed in only 3 of 2,509 cases Fig. 1. The *Oncotree Code* variable exhibited substantial class imbalance, with IMMC, PBS, BREAST, and MBC represented by very few samples, Fig. 2; these rare categories were therefore merged into a single group to reduce sparsity and improve model stability. Additionally, multiple variables contained many missing values (>500 entries). Rows with missing values in the *Chemotherapy* variable were removed because chemotherapy status is implicitly linked to several clinical and treatment-related features, and its absence often coincided with broader record incompleteness. Using *Chemotherapy* as the filtering criterion allowed for efficient removal of highly incomplete observations while retaining the clinical coherence of the dataset. In the final preprocessing step, survival-related outcome variables—*Overall Survival (Months), Overall Survival Status, Patient’s Vital Status, Relapse Free Status (Months)*, and *Relapse Free Status*—were excluded from the feature set. These variables represent post-diagnosis outcomes and were removed to prevent information leakage and ensure that subsequent analyses relied solely on baseline clinical and molecular features.

**Fig. 1:**
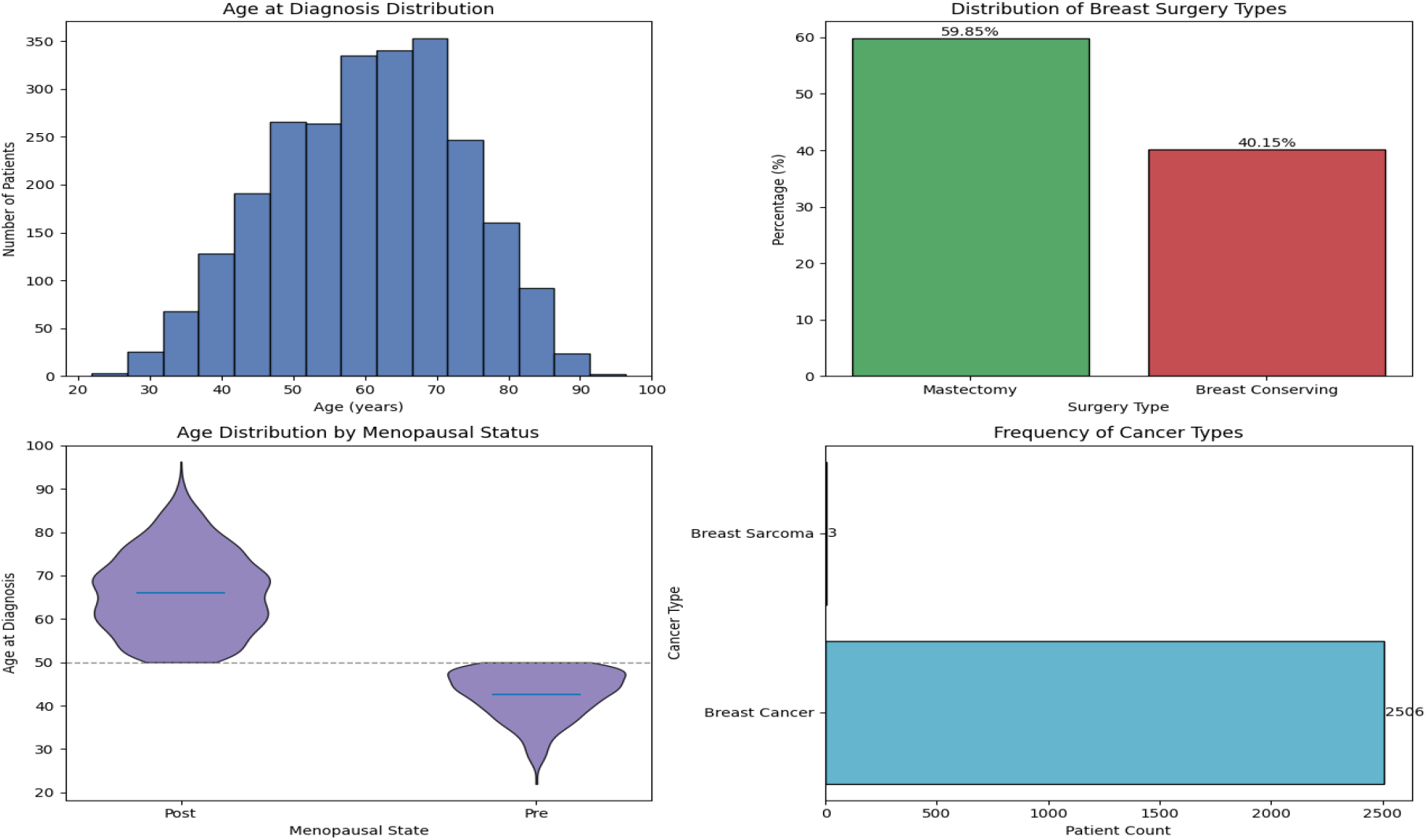
Distributions of age at diagnosis, breast surgery type, menopausal status by age, and cancer type in the METABRIC dataset.

**Fig. 2:**
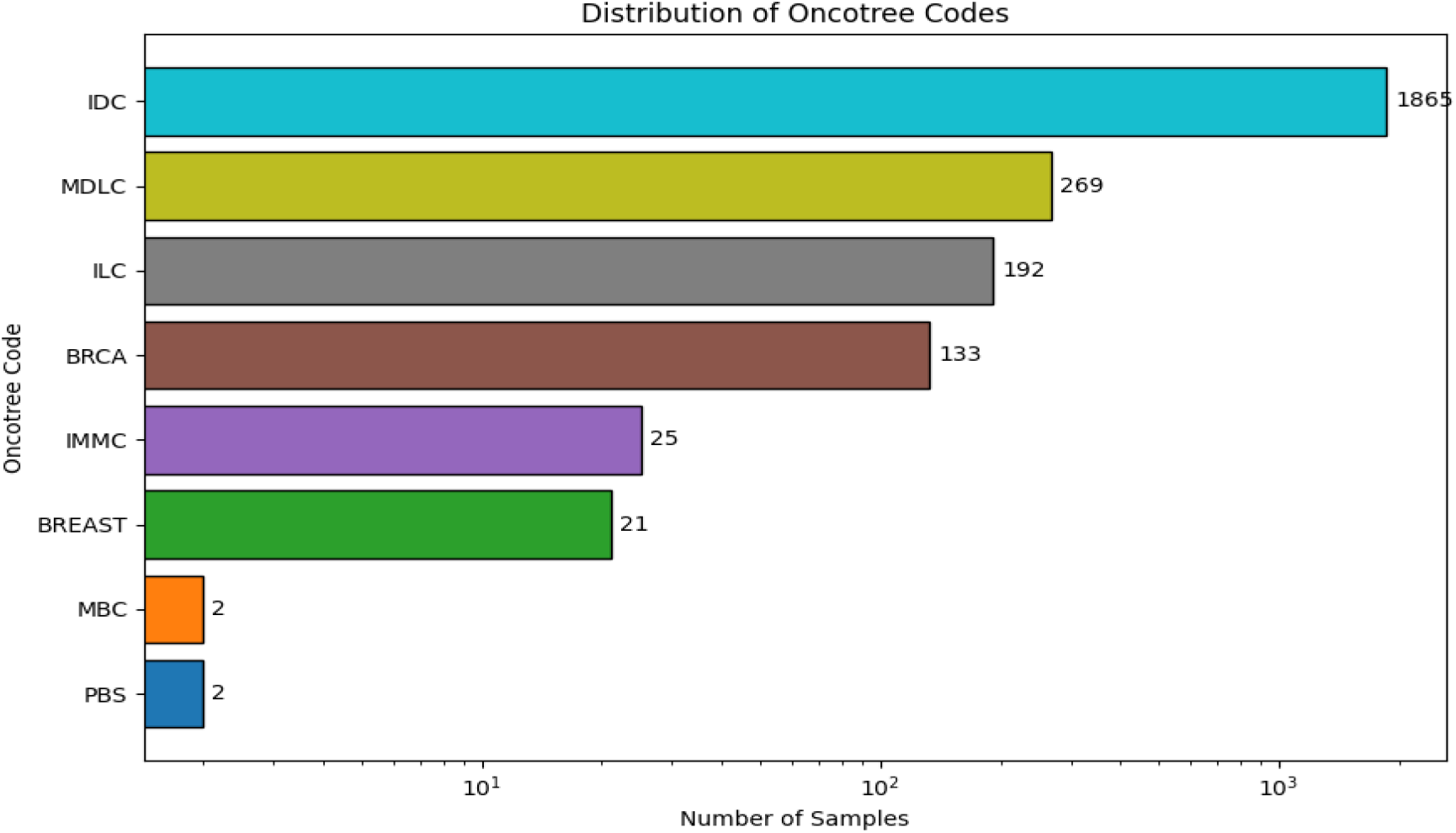
Distribution of tumor types by Oncotree code in the METABRIC dataset

Missing values present in the gene expression and CNV modalities were addressed using a weighted nearest neighbor imputation strategy [34]. Following established preprocessing protocols, gene expression measurements were discretized into three categories representing under-expression (−1), baseline expression (0), and over-expression (1), as described by Sun *et al*. and aligned with the methodology proposed by Gevaert *et al*. [35]. CNV data were encoded using five discrete states: −2, −1, 0, 1, and 2, reflecting varying degrees of copy number loss and gain. Clinical variables were normalized to the [0,1] range using min–max scaling, as defined in Equation (1), to ensure consistency across features and compatibility with downstream learning models [36].

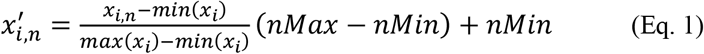

### Feature selection

Due to the high-dimensional, low-sample-size (HDLSS) characteristics of the dataset, direct application of deep learning models may result in limited efficiency and poor generalization [37]. The proposed framework integrates multimodal data, including CNV, gene expression, and clinical features. The CNV and gene expression modalities contain approximately 26,000 and 24,000 features, respectively, while the clinical modality comprises 22 patient-level variables such as age at diagnosis, cellularity, tumor size, and lymph node involvement, as described in the referenced dataset^1^. To address the HDLSS challenge, dimensionality reduction was performed using the minimum Redundancy Maximum Relevance (mRMR) algorithm [38, 39]. Feature selection was conducted incrementally, with features ranked by relevance and redundancy and evaluated using area under the ROC curve (AUC) across multiple cohorts. The number of selected features was progressively increased to identify an optimal balance between model complexity and predictive performance. Based on this procedure, 200 CNV features, 400 gene expression features, and 25 clinical variables were retained for the heterogeneous stacked random forest model. A detailed description of the selected features is provided in Table 2.

**Table 2:**
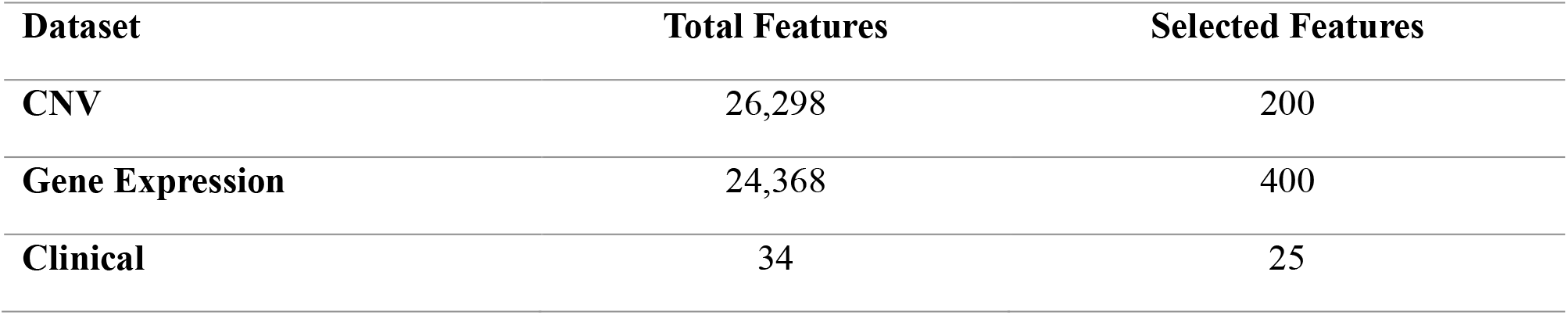
Feature selection detail.

| Dataset | Total Features | Selected Features |
| --- | --- | --- |
| CNV | 26,298 | 200 |
| Gene Expression | 24,368 | 400 |
| Clinical | 34 | 25 |

### A CNN and DNN-Based prediction for a UNI-MODAL dataset

Clinical and gene expression data are processed using CNN encoders, while CNV data are modeled with a DNN encoder, reflecting the structural differences among modalities. Each encoder operates independently in the first stage to learn modality-specific latent representations that are subsequently used for downstream prediction. The CNN architecture consists of a single one-dimensional convolutional layer with a stride of 2 and appropriate padding to control feature dimensionality, followed by a flattening operation and a fully connected layer with 150 hidden units. DNN comprises multiple fully connected layers with progressively reduced dimensionality, culminating in a 150-dimensional embedding. All layers employ the hyperbolic tangent (tanh) activation function. Network weights are initialized to a constant value of 0.1 for both CNN and DNN components. Regularization is applied to mitigate overfitting: L2 regularization is used in the CNN, while dropout with a rate of 0.5 is applied after each dense layer in the DNN. Model training is performed using the Adam optimizer due to its computational efficiency and favorable convergence properties. Since the task is binary classification, binary cross-entropy loss is employed to optimize the model. Model performance is evaluated using the area under the ROC curve (AUC). Experiments with varying batch sizes (8–128) indicate that a batch size of 8 achieves the best predictive performance and is therefore used in the final model configuration. Detailed architectural and parameter settings are provided in Fig. 3. Detailed tune parameters for CNN and DNN network is presented in Table 3.

**Table 3:**
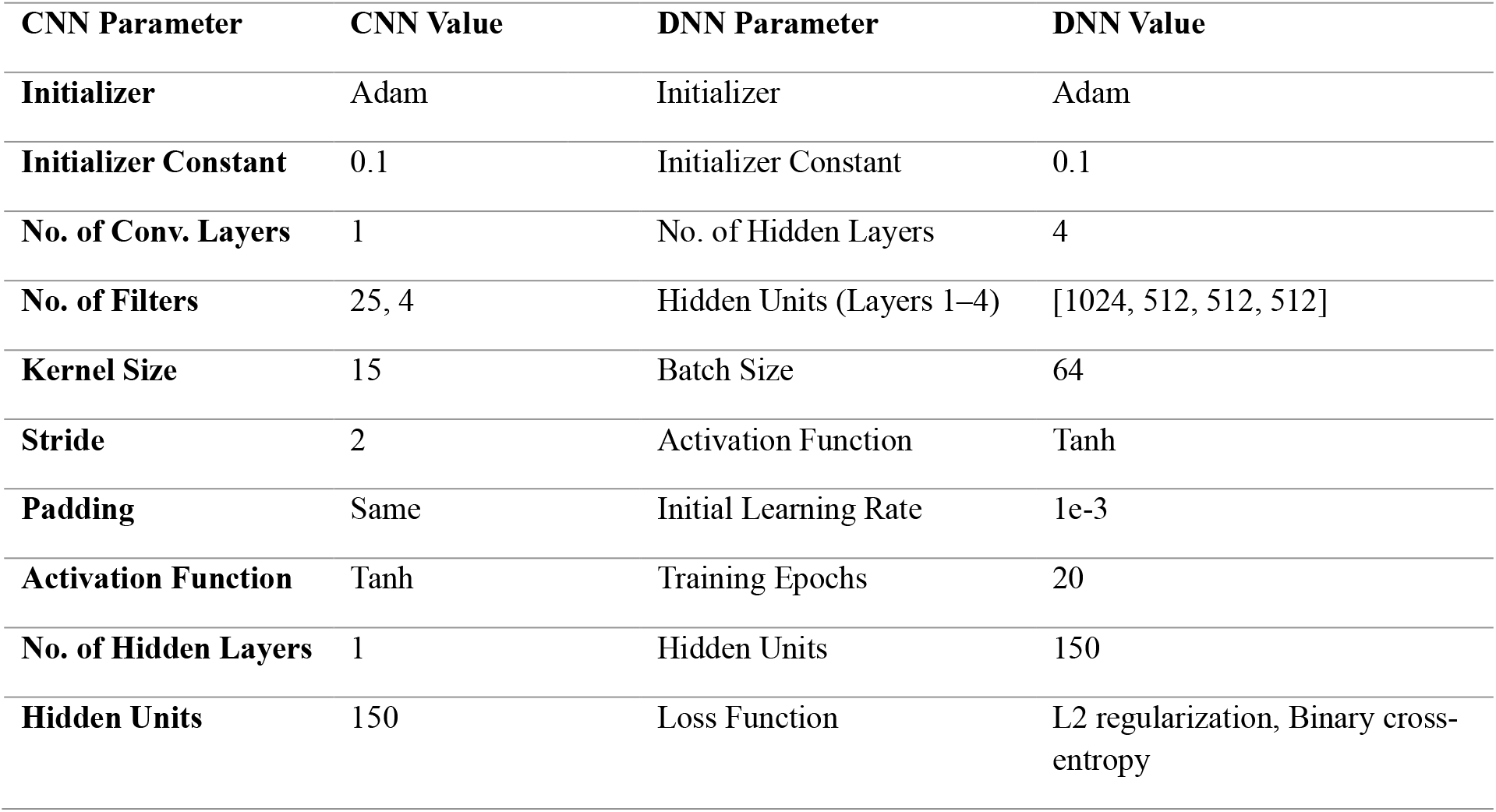
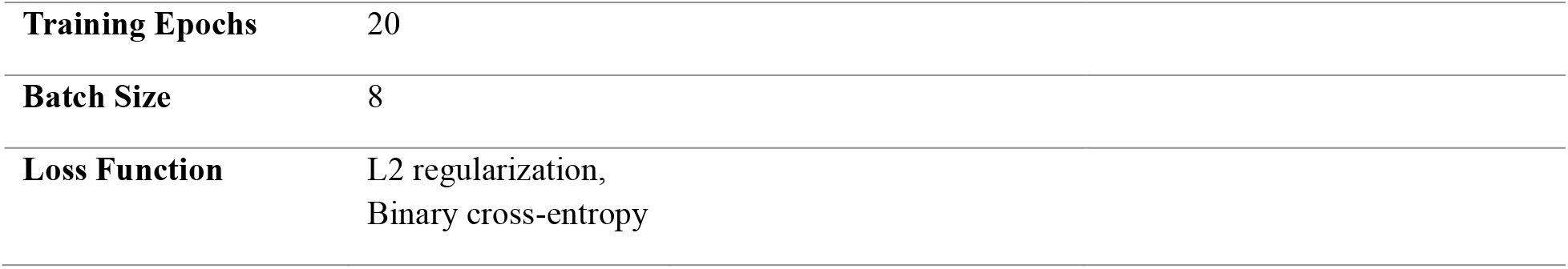
Summary of tuned hyperparameters for the unimodal CNN (clinical and gene expression) and DNN (copy number variation).

| CNN Parameter | CNN Value | DNN Parameter | DNN Value |
| --- | --- | --- | --- |
| Initializer | Adam | Initializer | Adam |
| Initializer Constant | 0.1 | Initializer Constant | 0.1 |
| No. of Conv. Layers | 1 | No. of Hidden Layers | 4 |
| No. of Filters | 25, 4 | Hidden Units (Layers 1–4) | [1024, 512, 512, 512] |
| Kernel Size | 15 | Batch Size | 64 |
| Stride | 2 | Activation Function | Tanh |
| Padding | Same | Initial Learning Rate | 1e-3 |
| Activation Function | Tanh | Training Epochs | 20 |
| No. of Hidden Layers | 1 | Hidden Units | 150 |
| Hidden Units | 150 | Loss Function | L2 regularization, Binary cross-entropy |

|  |  |
| --- | --- |
| **Training Epochs** | 20 |
| **Batch Size** | 8 |
| **Loss Function** | L2 regularization,<br>Binary cross-entropy |

**Fig. 3:**
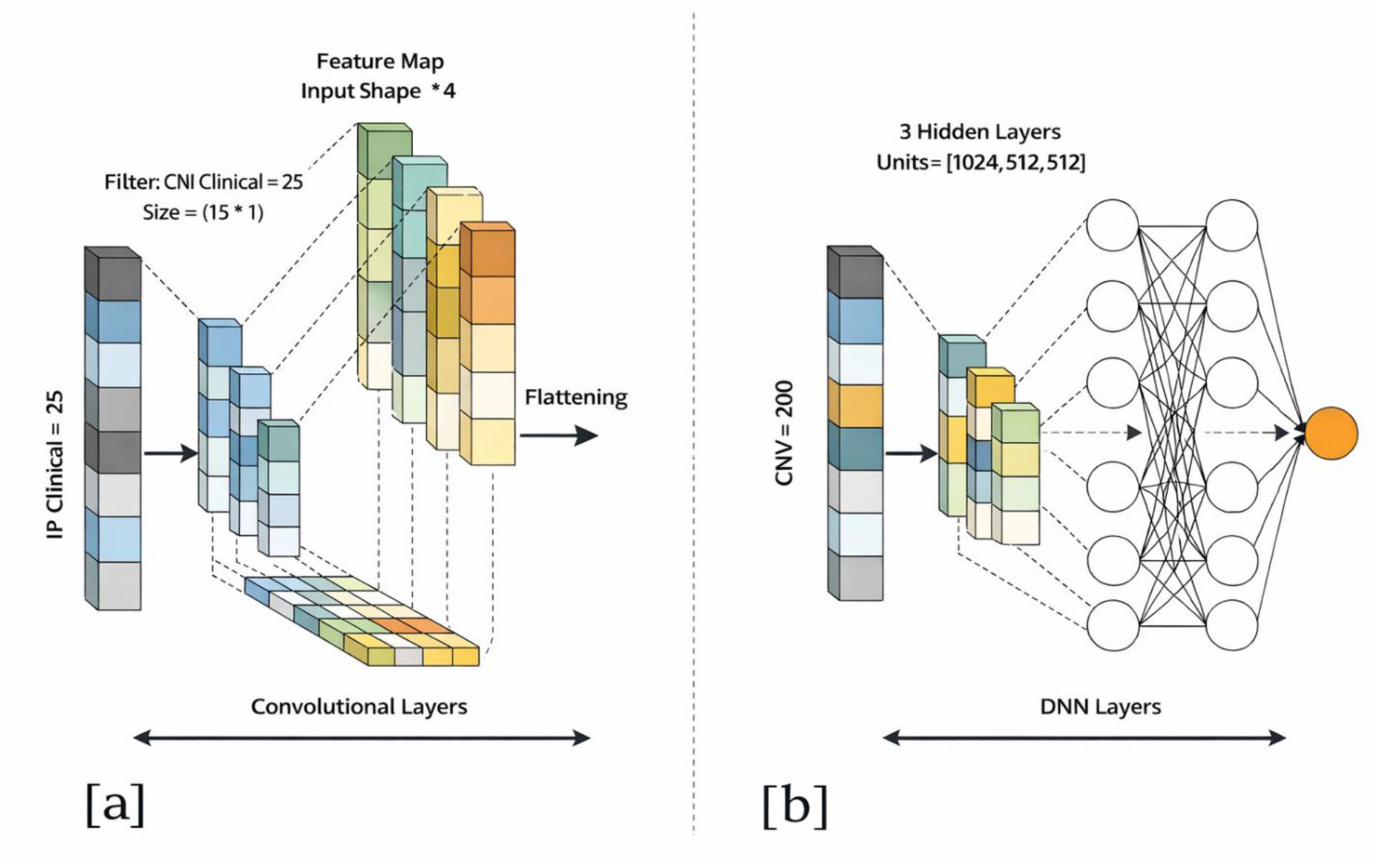
Network structures employed for unimodal representation learning: (a) CNN and (b) DNN.

**Fig. 4:**
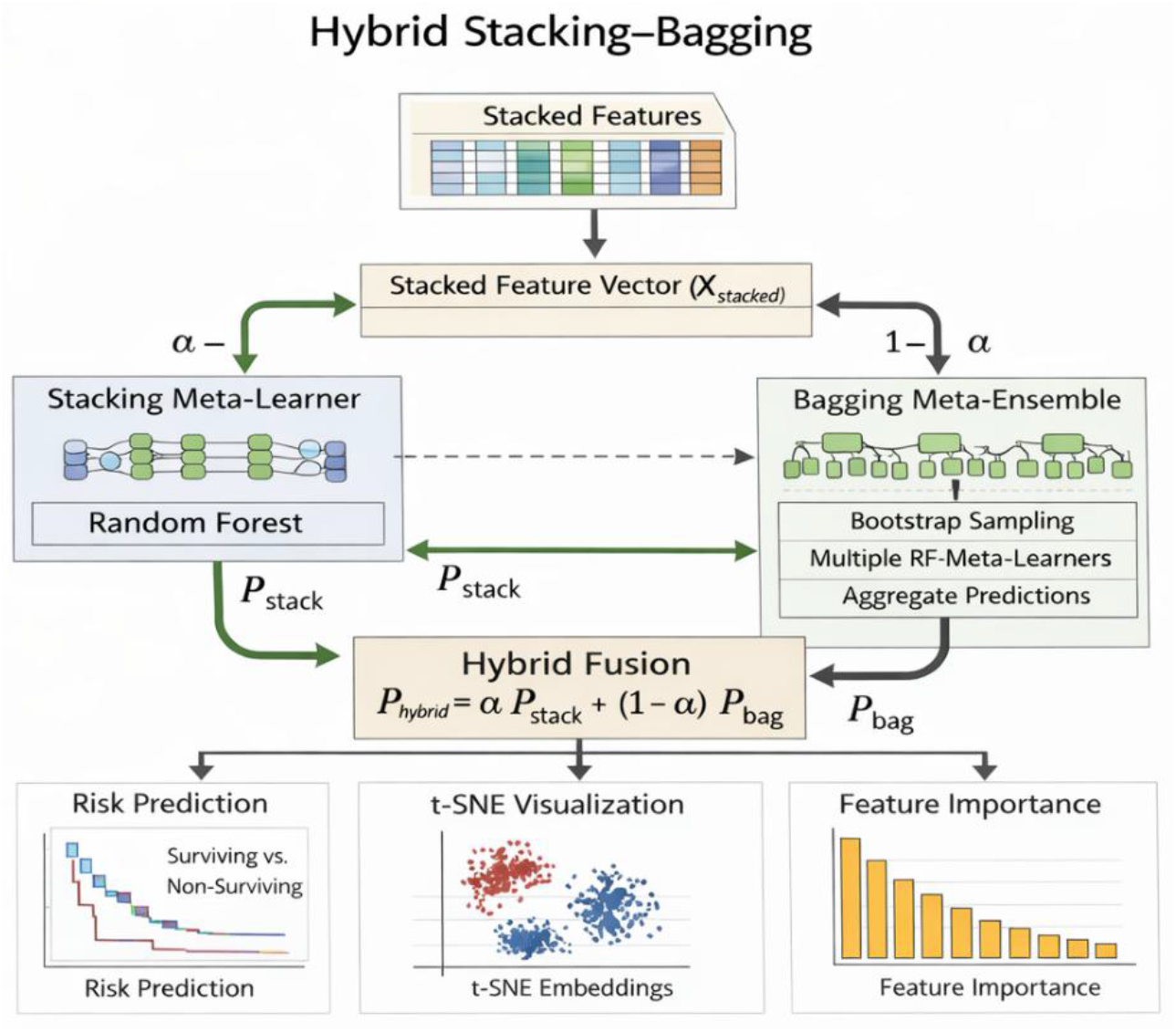
The systematic proposed framework.

**Fig. 5:**
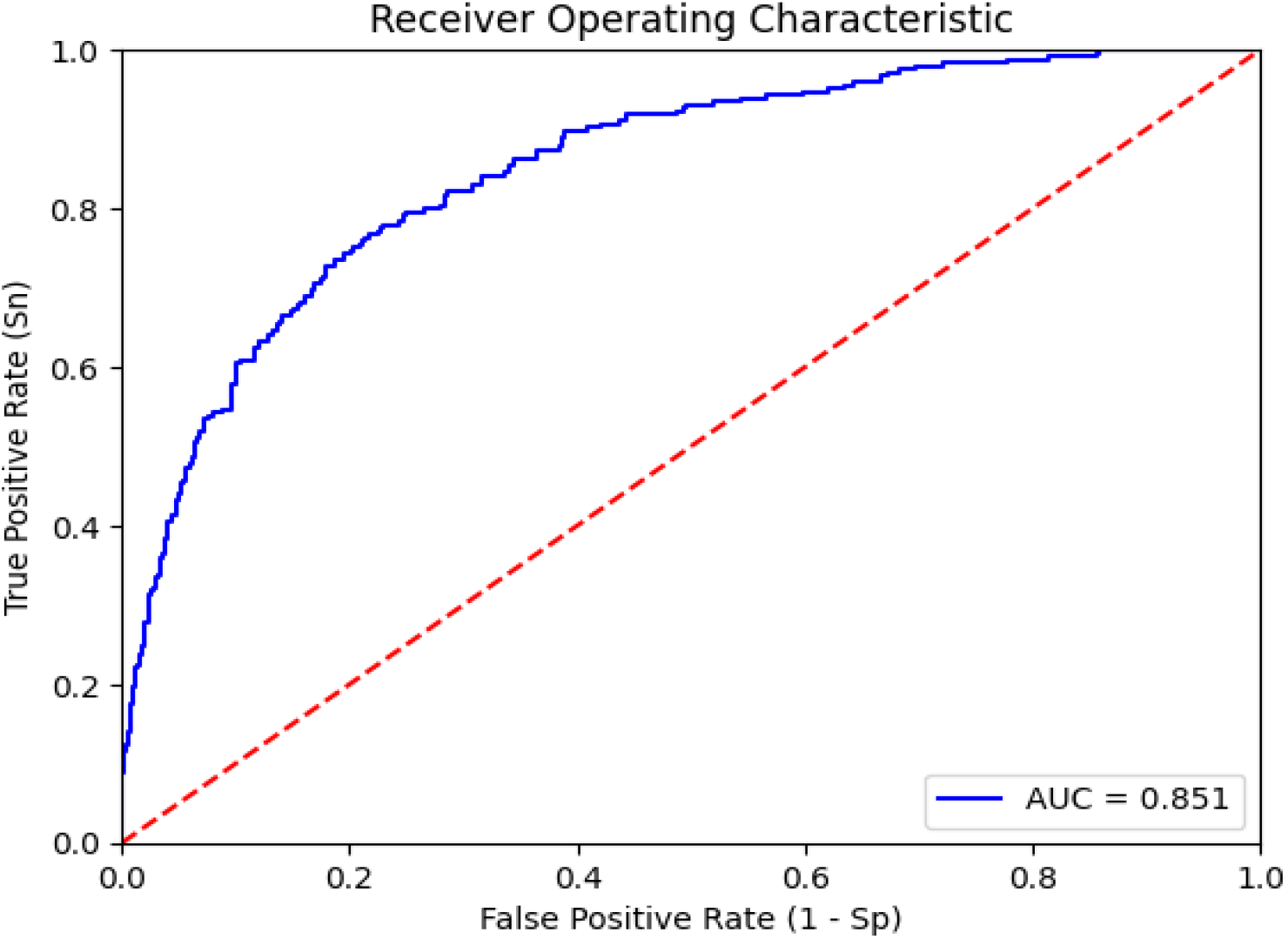
Clinical unimodal

**Fig. 6:**
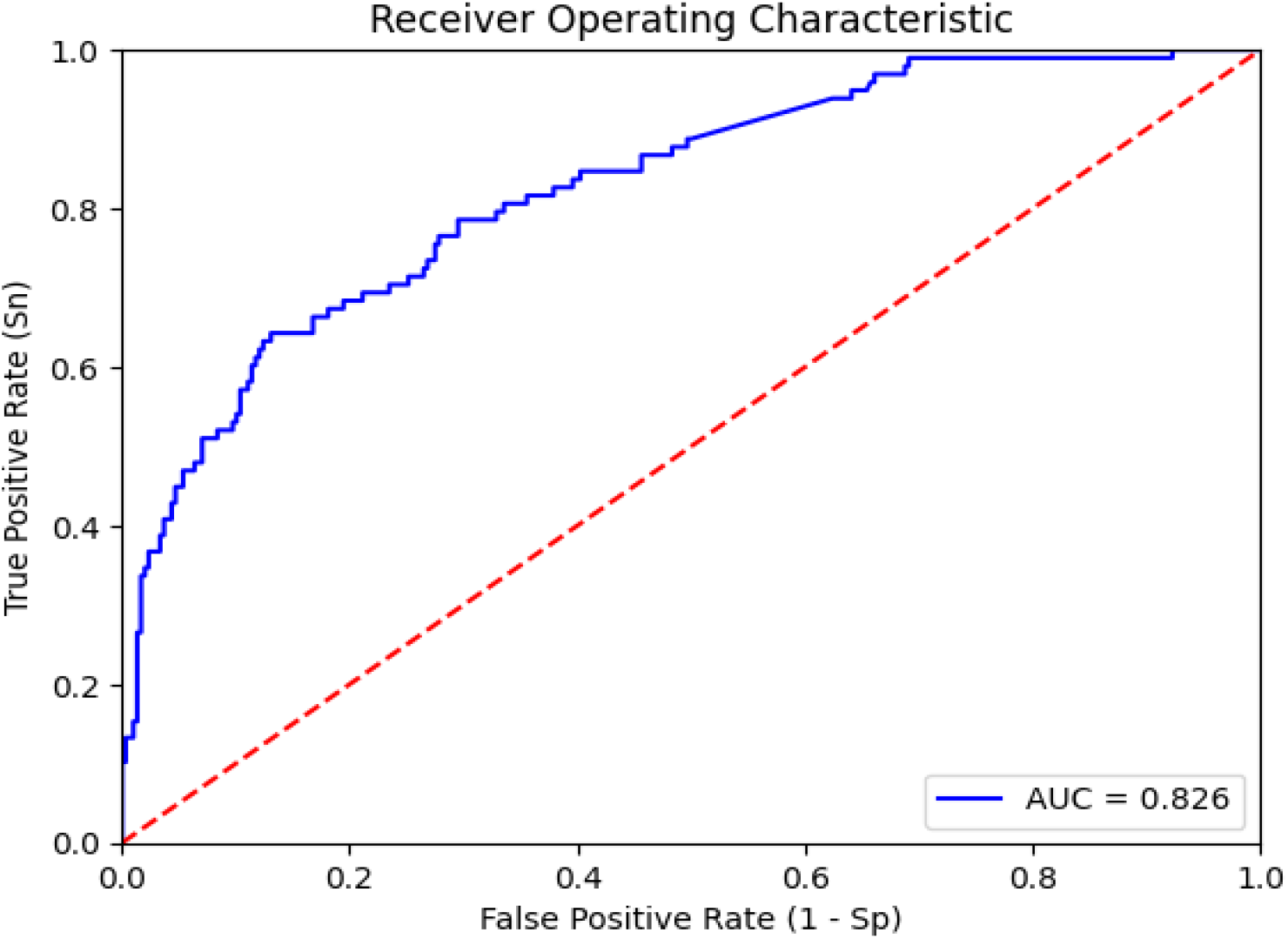
CNV unimodal

**Fig. 7:**
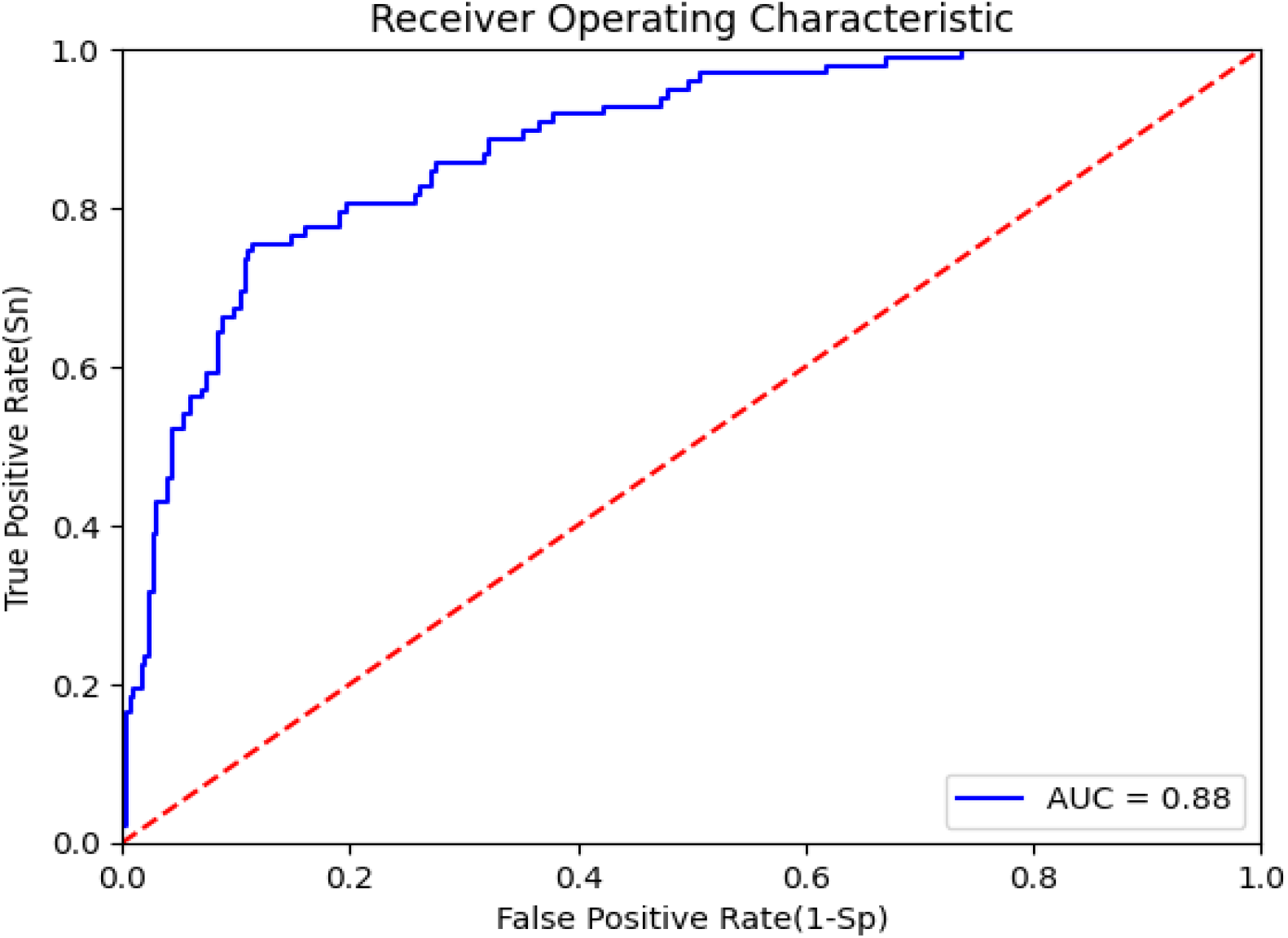
gene expression

**Fig. 8:**
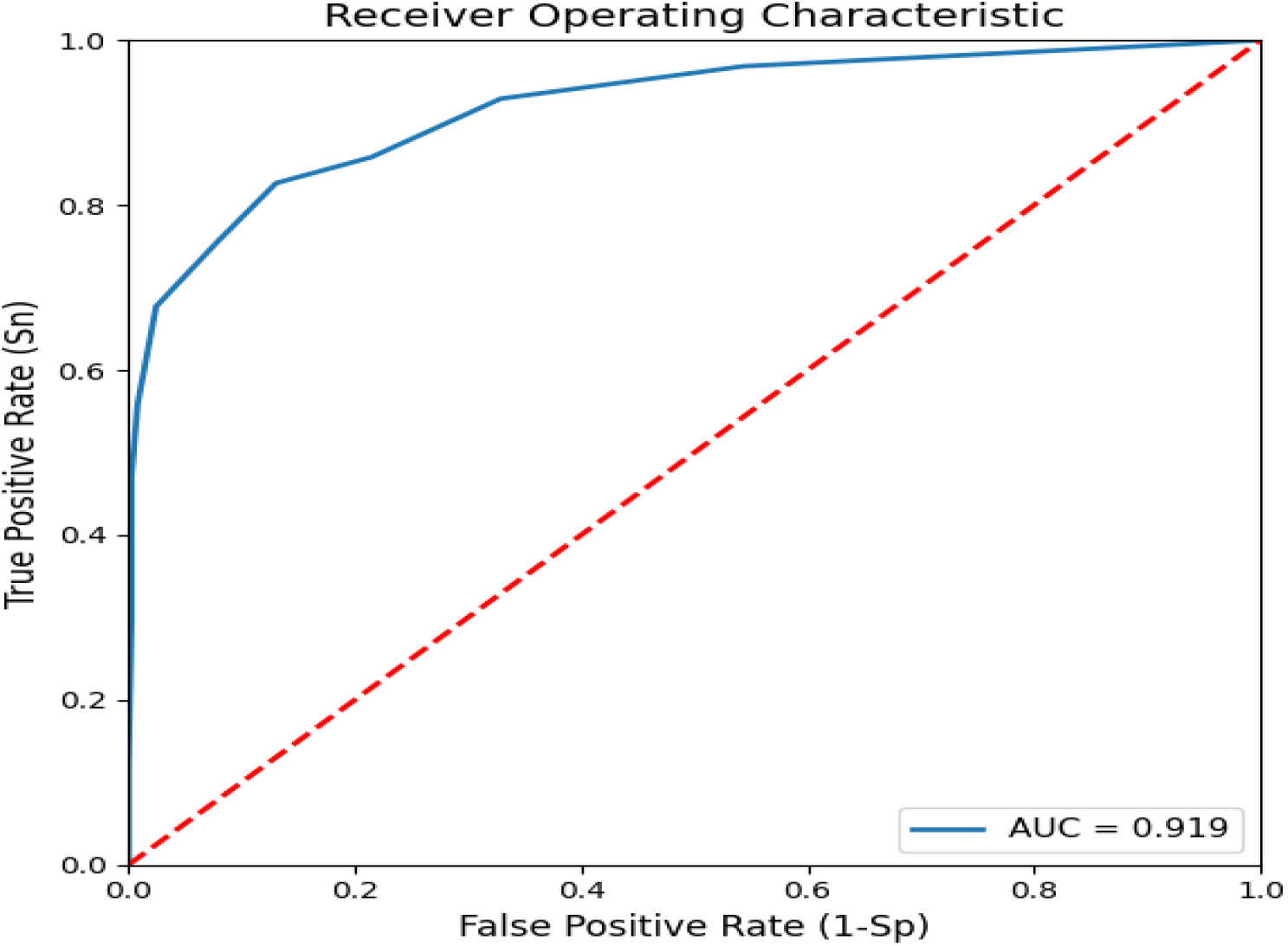
Ensemble Stacking

**Fig. 9:**
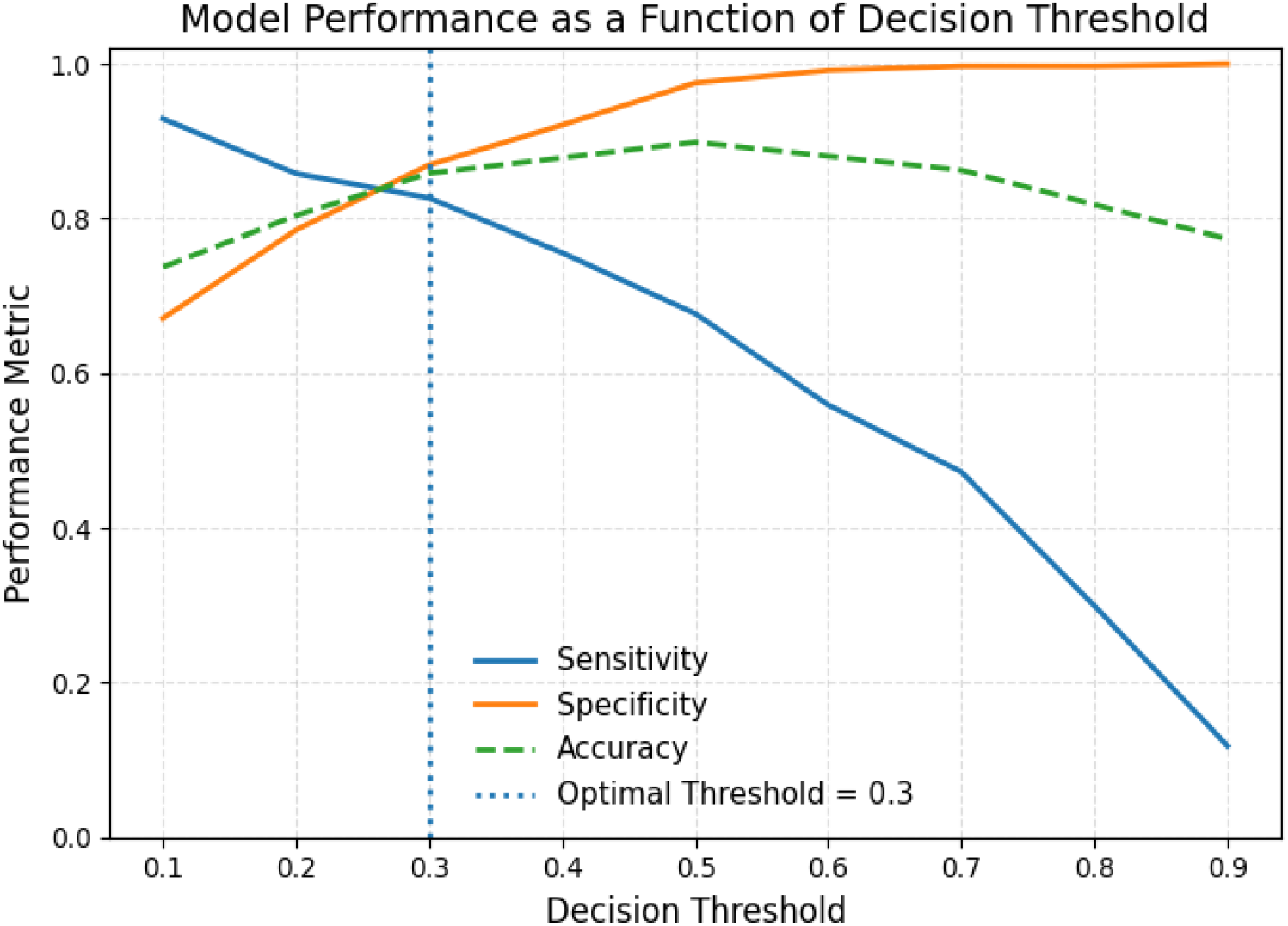
Trade-off between sensitivity, specificity, and accuracy across decision thresholds, highlighting the optimal operating point.

**Fig. 10:**
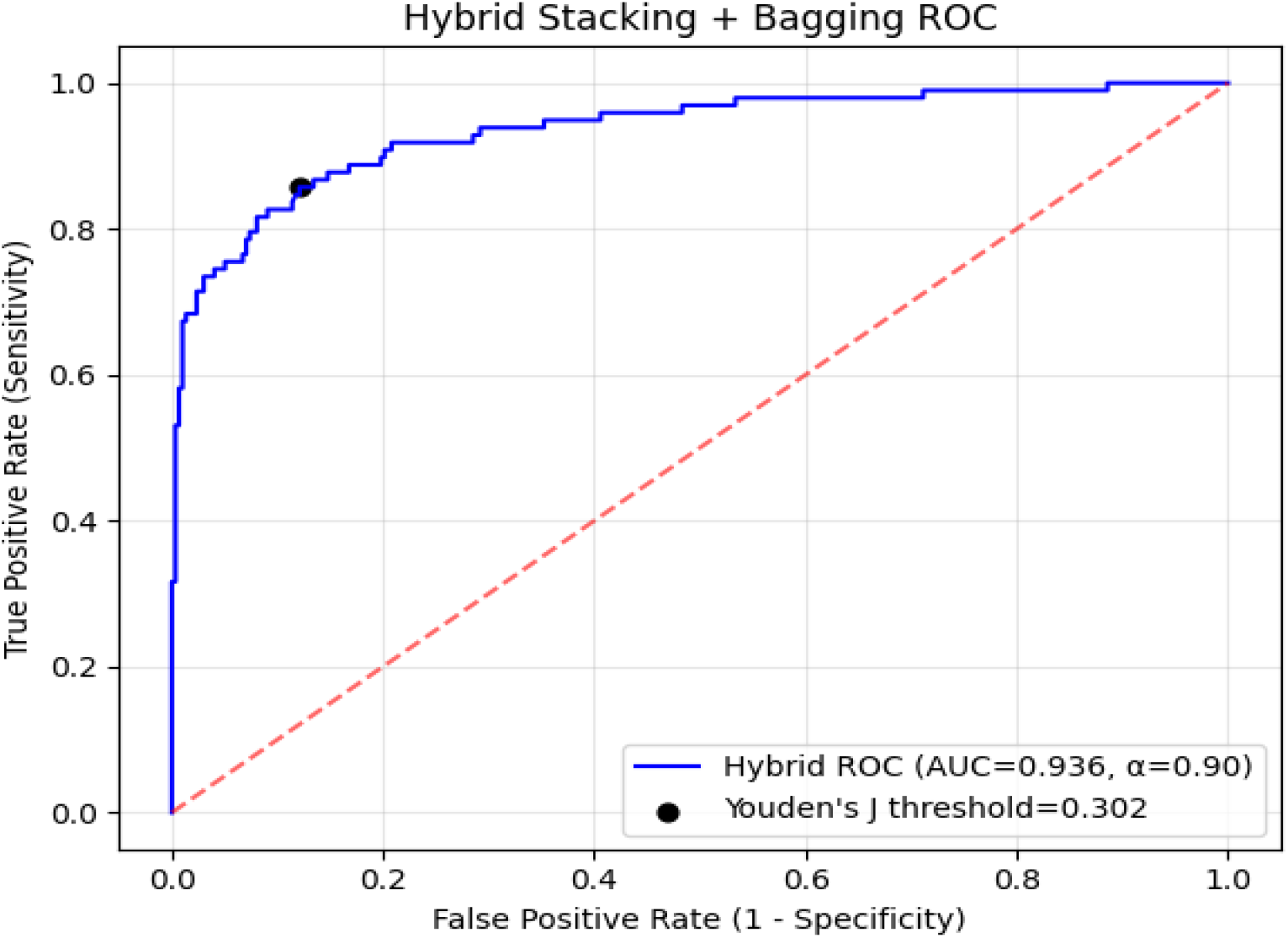
Hybrid Model

**Fig. 11:**
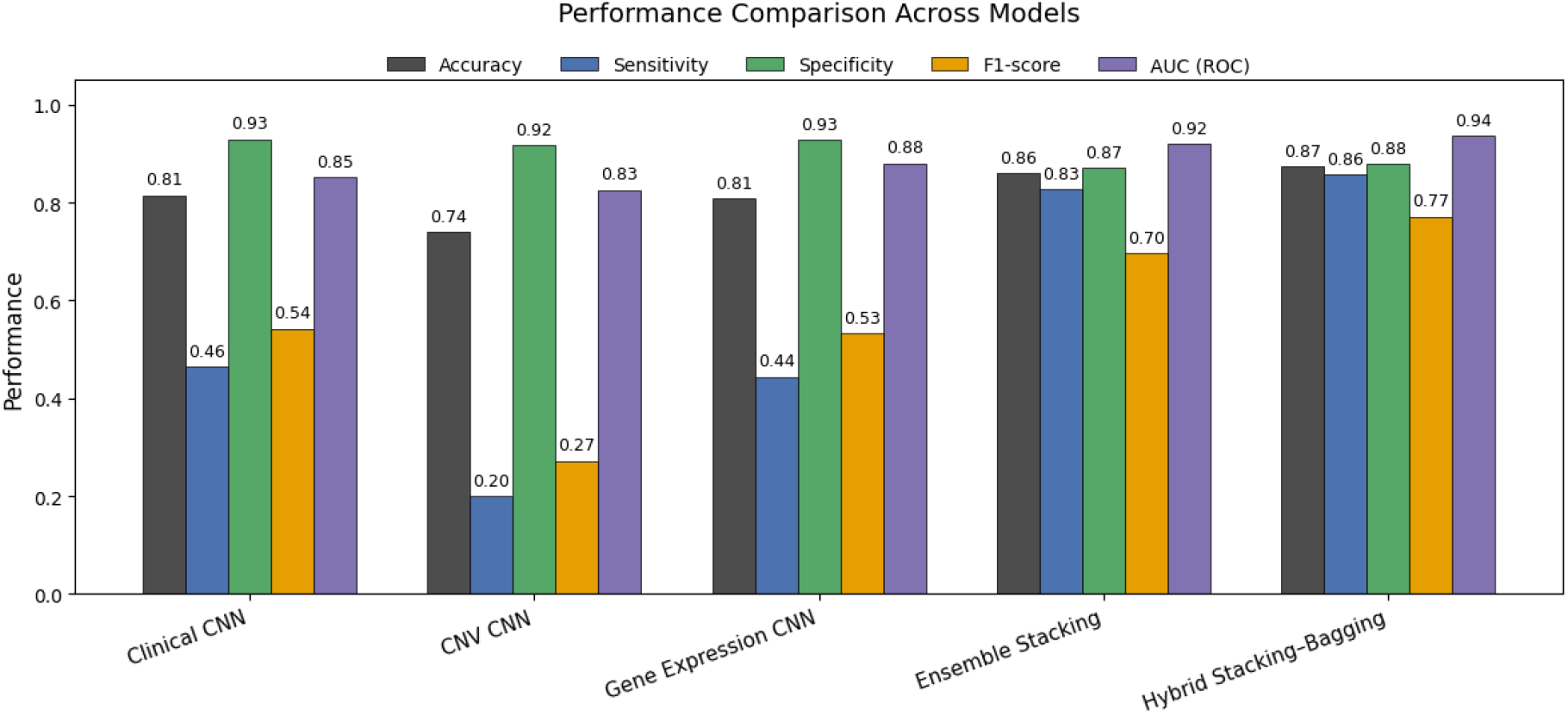
Performance comparison of unimodal models, ensemble stacking, and the proposed hybrid stacking–bagging framework.

### Proposed Model setup

The proposed method introduces a heterogeneous multimodal hybrid ensemble framework that combines stacking and bagging to improve robustness and predictive accuracy in breast cancer risk classification. The framework integrates clinical features, gene expression profiles, and copy number variation (CNV) data within a unified ensemble architecture designed to reduce variance and mitigate overfitting commonly observed in conventional ensemble models. For each patient, features derived from the clinical, gene expression, and CNV modalities are first combined into a single stacked feature representation. This stacked feature vector serves as a shared input for both ensemble branches of the framework, enabling joint learning across heterogeneous data sources while preserving complementary modality-specific information. The stacking component employs a Random Forest meta-learner trained on the stacked feature vector to capture nonlinear relationships and interactions among multimodal features. This meta-learner produces a probabilistic output *P*_stack_, representing the risk prediction obtained through stacked multimodal fusion. In parallel, a bagging-based meta-ensemble is constructed using bootstrap sampling of the stacked feature vector. Multiple Random Forest meta-learners are trained on different bootstrap samples, and their predictions are aggregated to generate a bagging-based probability output *P*_bag_, thereby improving prediction stability and reducing model variance. To leverage the complementary strengths of stacking and bagging, the final risk prediction is obtained through a weighted hybrid fusion of the two probabilistic outputs. Specifically, the hybrid prediction is computed as

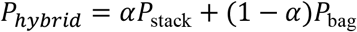

where *α*controls the relative contribution of the stacking and bagging components. The fused probability *P*_hybrid_is subsequently used to generate the final breast cancer risk classification.

Beyond classification, the framework supports interpretability and visualization. Feature importance scores derived from the Random Forest meta-learners provide insights into the contribution of clinical and molecular features, while t-SNE embeddings of the stacked feature representation are used to visualize class separability and assess the discriminative structure learned by the hybrid ensemble.

## Results and Discussion

To establish a baseline for evaluating multimodal integration, we first assessed the predictive performance of unimodal models trained independently on clinical features, gene expression profiles, and copy number variation (CNV) data from the METABRIC cohort. This step provides a reference point for understanding the individual contribution and discriminative capacity of each data modality and highlights the limitations of relying on a single source of information for breast cancer risk prediction. The unimodal results also serve as a benchmark against which the performance gains achieved through stacking and hybrid ensemble strategies can be systematically compared.

Building on the unimodal baseline results, we next evaluated a multimodal stacking ensemble to examine whether joint learning across clinical, gene expression, and CNV data could improve predictive performance. Unlike unimodal models, the stacking approach integrates complementary information from multiple data sources through a meta-learning strategy, enabling more effective capture of complex biological interactions. This comparison allows us to quantify the performance gains achieved through multimodal fusion alone, prior to introducing additional variance reduction via the proposed hybrid stacking–bagging framework.

To determine the optimal decision threshold, model performance was evaluated across a range of thresholds using accuracy, sensitivity, and specificity (Table 4). Rather than selecting a threshold based solely on accuracy, which can be misleading in the presence of class imbalance, the optimal threshold was identified by maximizing Youden’s index, defined as sensitivity + specificity − 1. This criterion provides a balanced measure of diagnostic effectiveness by simultaneously accounting for both false positive and false negative rates. As shown in Table 4, a decision threshold of 0.3 yielded the maximum Youden’s index, indicating the most favorable trade-off between sensitivity and specificity. At this threshold, sensitivity and specificity are well balanced, and overall classification accuracy is near its maximum. Thresholds lower than 0.3 increase sensitivity at the expense of substantially reduced specificity, while higher thresholds improve specificity but lead to a pronounced loss in sensitivity. Therefore, the threshold of 0.3 was selected as the optimal operating point for subsequent analyses.

**Table 4:** Sensitivity, specificity, accuracy, and Youden’s index evaluated across decision thresholds.

| Threshold | Sensitivity | Specificity | Youden |
| --- | --- | --- | --- |
| **0.2** | ~0.86 | ~0.79 | ~0.65 |
| **0.3** | **~0.83** | **~0.86** | **~0.69 ← max** |
| **0.4** | ~0.75 | ~0.92 | ~0.67 |
| **0.5** | ~0.68 | ~0.98 | ~0.66 |

While the stacking ensemble demonstrates clear performance gains over unimodal baselines, its predictive performance can still be influenced by variance arising from the meta-learning process. To further enhance stability and generalization, we next evaluated the proposed hybrid stacking–bagging framework, which combines multimodal stacking with bootstrap-based aggregation. This final experiment assesses whether incorporating bagging into the stacking architecture provides additional improvements in discrimination performance and more balanced sensitivity–specificity trade-offs compared to stacking alone.

## Conclusion

This study presented a heterogeneous multimodal hybrid ensemble framework that integrates stacking and bagging to improve breast cancer risk prediction using clinical features, gene expression profiles, and copy number variation data. By systematically evaluating unimodal models, ensemble stacking, and the proposed hybrid approach, we demonstrated that combining complementary data modalities within a robust ensemble architecture leads to substantial performance gains over single-modality and conventional stacking methods. The unimodal results confirmed that no individual data source is sufficient to capture the complex biological mechanisms underlying breast cancer outcomes, with gene expression and clinical data providing moderate discriminative power and CNV data showing comparatively weaker performance. While ensemble stacking significantly improved predictive accuracy by leveraging multimodal integration, its performance remained sensitive to variance introduced at the meta-learning stage. The proposed hybrid stacking–bagging framework addressed this limitation by incorporating bootstrap-based aggregation, resulting in further improvements in ROC AUC, accuracy, and F1-score, as well as more balanced sensitivity and specificity at the optimal operating threshold.

Beyond improved classification performance, the hybrid framework offers practical advantages for clinical decision support. The use of ensemble-based meta-learners enables interpretability through feature importance analysis, while visualization of the learned feature space highlights enhanced class separability achieved through hybrid fusion. Importantly, the proposed approach is scalable to high-dimensional multi-omics data and does not rely on modality-specific deep learning architectures, making it adaptable to a wide range of biomedical prediction tasks. Despite these promising results, this study has limitations. The analysis was conducted on a single cohort, and external validation on independent datasets will be necessary to assess generalizability. Additionally, future work could explore adaptive or data-driven strategies for optimizing the fusion weight between stacking and bagging components, as well as extending the framework to time-to-event survival modeling.

In summary, the proposed hybrid stacking–bagging ensemble provides a robust and effective strategy for multimodal breast cancer risk prediction, offering improved stability, accuracy, and interpretability. This work highlights the value of hybrid ensemble learning for integrating heterogeneous biomedical data and represents a meaningful step toward more reliable precision oncology models.

## Notes

### Competing Interest Statement

The authors have declared no competing interest.

